# Alkaline ceramidase catalyzes the hydrolysis of ceramides via a catalytic mechanism shared by Zn^2+^-dependent amidases

**DOI:** 10.1101/442202

**Authors:** Jae Kyo Yi, Ruijuan Xu, Lina M. Obeid, Yusuf A. Hannun, Michael V. Airola, Cungui Mao

## Abstract

Human alkaline ceramidase 3 (ACER3) is one of three alkaline ceramidases (ACERs) that catalyze the conversion of ceramide to sphingosine. ACERs are the members of the CREST superfamily of integral-membrane lipid hydrolases, including the adiponectin receptors which play roles in energy metabolism. All CREST members conserve a set of three Histidine, one Aspartate, and one Serine residue. However, the structural and catalytic roles for these residues are unclear. Here, we use ACER3 as a prototype enzyme to gain insight into this unique class of enzymes. Recombinant ACER3 was expressed in yeast cells that lack endogenous ceramidase activity, and microsomes were used for biochemical characterization. Six point mutantions of the conserved CREST motif were developed that are predicted to form a Zn-dependent active site based on homology with the human adiponectin receptors, whose crystal structures were recently determined. Five mutations completely lost their activity, except for S77A, which showed a 600-fold decrease compared with the wild-type enzyme. The activity of S77C mutation was pH sensitive, with neutral pH partially recovering ACER3 activity. This suggested a role for S77 in stabilizing the oxyanion of the transition state and differs from the proposed role in Zinc coordination for the adiponectin receptors (Vasiliauskaité-Brooks et. al., Nature, 2017). Together, these data suggest ACER3 is a Zn2+-dependent amidase that uses a catalytic mechanism for ceramide hydrolysis that is similar to other soluble Zn-based amidases. Consistent with this mechanism, ACER3 was specifically inhibited by trichostatin A, an HDAC inhibitor, which is a strong chelator of Zinc.

## INTRODUCTION

Ceramidases catalyze the hydrolysis of ceramides to release free fatty acids and sphingosine (SPH). As such, they are important intermediates of complex sphingolipids that play an important role in the integrity and function of cell membranes ^1^. Alkaline ceramidase 3 (ACER3) is one of three ACERs (ACER1, 2, and 3), which were cloned by our group ^2^. ACER3 is localized to both the Golgi complex and endoplasmic reticulum (ER), and it is highly expressed in various tissues compared with the other two alkaline ceramidases. Additionally, it was demonstrated that ACER3 hydrolyzes a synthetic ceramide analogue NBD-C_12_-phytoceramide *in vitro*; thus, it was formerly named the human alkaline phytoceramidase (haPHC) ^2^. Later, we reported that ACER3 catalyzes the hydrolysis of ceramides, dihydroceramides, and phytoceramides carrying an unsaturated long acyl chain (C18:1 or C20:1) with similar efficiency ^3^. Recently, important roles of ACER3 in neurobiology have been highlighted. We demonstrated that ACER3 deficiency results in Purkinje cell degeneration in mice ^4^ and that a point mutation of ACER3 leads to progressive leukodystrophy in early childhood in humans ^5^.

The sequence similarity between ACERs, and progestin and adipoQ receptors (PAQR receptors) was revealed by Villa *et al.* ^6^. In turn, ACER3 has been reported to belong to a large superfamily of proteins referred to as CREST (for alkaline <u>c</u>eramidase, PAQR receptor, P<u>e</u>r1, <u>S</u>ID-1 and <u>T</u>MEM8) ^7^. The CREST superfamily is highlighted in various cellular functions and biochemical activities. Ceramidases are lipid-modifying enzymes ^8^. The PAQR receptors contain the adiponectin receptors, which regulate energy metabolism ^9–11^. The Per1 family consists of fatty acid remodeling hydrolases for GPI-anchored proteins ^12–13^. SID-1 family of putative RNA transporters is involved in systematic RNA interference ^14–15^. Lastly, the TMEM8 family is known to play a role in cancer biology as putative tumor suppressors ^16–17^.

All members of the CREST superfamily conserve a set of three Histidine residues, a Serine residue, and an Aspartate residue. These residues are the defining motif of the CREST family and were suggested to form a metal-dependent active site for lipid hydrolysis. However, the specific roles of the conserved residues in CREST proteins have not been reported. The independent discoveries of hydrolase activities in alkaline ceramidases ^2, 18^ and Per1 ^13^ suggest that the majority of CREST members are metal-dependent hydrolases. In support of this, the adiponectin receptors were recently found to also hydrolyze ceramide, but at a slow rate ^19^.

The crystal structures of the human adiponectin receptors (ADPR1 and ADPR2), which represent the first structures for any CREST member, were recently determined ^20^, although there was a limitation of the study due to lack of presenting a direct catalytic approach. ADPR1 and ADPR2 exhibit a seven-transmembrane helix architecture with a zinc ion located near the end of a large internal cavity as the putative active site. Importantly, the zinc ion is coordinated by the three His residues of the CREST motif, and the conserved Ser and Asp residues are in close proximity to the Zinc ion. The His residues are required for activation of the adiponectin-stimulated pathways.

A catalytic mechanism for the ADPRs was recently proposed based on molecular dynamics simulations ^19^. In this mechanism, the zinc coordinating residues were proposed to rearrange to facilitate ceramide hydrolysis. This mechanism differs quite drastically from other similar Zn-based amidases, such as HDACs, LpxC, peptide deformylase, matrix metalloproteinases (MMPs), and human neutral ceramidase ^21–24^. In all of these cases, the zinc coordinating residues do not rearrange and an Asp or His residue serves as a general base for catalysis.

Compared with the biological role of ACER3, which has been studied in the last decade, the kinetic mechanism of its intrinsic ceramidase activity has not been described in detail. Thus, there is little understanding of the catalytic properties of ACER3. In this study, our biochemical results and mutational analysis allow us to propose the mechanism for how ACER3 hydrolyzes ceramides, which most likely applies to other CREST members. Furthermore, the inhibitor assay exhibits a possibility to develop direct and specific inhibitors of ACER3 and its related paralogs, ACER1 and ACER2, for treatments of diseases associated with dysregulation of the metabolism of ceramides and other sphingolipids, such as cancers, diabetes mellitus, cardiovascular diseases, neurodegenerative diseases, and skin diseases ^4–5, 25–26^.

## EXPERIMENTAL PROCEDURES

### Site-directed mutagenesis

The yeast expression plasmid pYES2-ACER3 that contains the FLAG-tagged open reading frame (ORF) of the wild-type *ACER3* gene was constructed in our previous study ^2, 18^ and used as a template for site-directed mutagenesis. The codon of His81, His217, His22, Asp92, Ser77 in the ACER3 ORF in pYES2-ACER3 (WT) was switched to a codon of Ala or Cys using QuickChange II XL a Site-Directed Mutagenesis Kit (Agilent Technology; Danbury, CT) following the manufacturer’s manual. The resulting plasmids containing the mutated ORF was sequenced to confirm the codon switch.

### Protein expression in yeast cells

The *Saccharomyces cerevisiae* mutant strain ∆*ypc1*∆*ydc1*, in which both yeast alkaline ceramidase genes *YPC1* and *YDC1* was deleted ^2^, was grown at 30°C on agar plates containing YPD medium (1% yeast extract, 2% peptone, 2% dextrose), or in liquid YPD medium with rotational shaking at 200 rpm. ∆*ypc1*∆*ydc1* cells were transformed with WT, H81A, H217A, H221A, D92A, S77A, and S77C, respectively, and the resulting transformants were selected on agar plates containing uracil-dropout synthetic medium (SC-Ura) with 2% glucose. ∆*ypc1*∆*ydc1* yeast cells transformed were grown in SC-Ura medium with 2% galactose to induce the expression of ACER3 or mutated ACER3s, respectively. Total membranes were prepared from yeast cells, and the expression of wild-type or mutanted ACER3s was determined by Western blot analyses using anti-FLAG antibody as described ^18^.

### Sample preparation and enzyme reaction for the ceramidase activity assays

The total membranes were collected from transformed yeast cells as described in ^18^. Briefly, cells were harvested in log phage (at OD ≅ 1.0) and washed by PBS. Cells were broken by acid-washed beads and transferred to ultracentrifugation tubes. Then, all cell membranes were pelleted by ultracentrifugation (at 100,000 g for 1 hr with a TLA55 rotor, Beckman Coulter Life Sciences, Brea, CA) from post-nuclear cell lysates in Buffer A (25 mM Tris-HCl, pH 7.4, 0.25 M sucrose) supplemented with a protease inhibitor mixture (Roche, Indianapolis, IN). Membrane pellets were resuspended in Buffer B (25 mM Tris, pH7.4, 5 mM CaCl2 and 150 mM NaCl) by brief sonication.

Membrane homogenates (1 μg of microsomal proteins) were measured for alkaline ceramidase activity using D-_ribo_-C_12_-NBD-phytoceramide (NBD-C_12_-PHC, Avanti, Alabaster, AL) as a substrate. Briefly, NBD-C_12_-PHC was dispersed by water bath sonication in Buffer C (25 mM glycine-NaOH, pH 9.4, 5 mM CaCl_2_, 150 mM NaCl, and 0.3% Triton X-100). The lipid-detergent mixtures were boiled for 30 s and chilled on ice immediately to form homogeneous lipid-detergent micelles, which were mixed with an equal volume of membranes suspended in Buffer B. Enzymatic reaction was carried out at 37 °C for 30 min, and the extraction buffer (chloroform: Methanol, 1:1) was added with 3 volumes of reaction mixture to quench the reactions. After centrifugation, the organic phase was collected and dried under a stream of inert nitrogen gas. Dried samples were dissolved in Mobile Phase B consisted of 1 mM ammonium formate in methanol containing 0.2% formic acid for high-performance liquid chromatography (HPLC).

### High-performance liquid chromatography (HPLC)

HPLC was utilized to measure ceramidase activities and kinetics of ACER3 with a modification of the technique described in ^27^. Ten μL of each reaction was injected into reverse phase LC and separated by a Spectra C8SR Column (150 × 3.0 mm; 3 μm particle size; Peeke Scientific, Redwood City, CA). Mobile Phase A consisted of 2 mM ammonium formate in water containing 0.2% formic acid. Mobile Phase B consisted of 1 mM ammonium formate in methanol containing 0.2% formic acid. Fluorescence was determined using an 1100 Series HPLC-FLD Fluorescent Detector (Agilent, Santa Clara, CA) set to excitation and emission wavelengths of 467 and 540 nm, respectively. Fluorescent peaks were identified by comparing their retention times with NBD-C_12_-FA and NBD-C_12_-PHC (Fig. S3). Calibration curves were calculated by linear regression. The data were exported to Excel spreadsheets to generate calibration lines and calculate sample concentrations.

### ACER3 inhibitor assay

Before enzymatic reaction was performed, collected microsomes were pre-treated with inhibitors including DMSO (control), Trichostatin A (TSA, Sigma-Aldrich, St. Louis, MO) or C_6_-Cer-Urea (Avanti, Alabaster, AL) for 30 min. The lipid-detergent mixtures were then mixed with an equal volume of microsomes. Following incubation at 37 °C for 30 min, the reaction was terminated by adding the extraction buffer (chloroform/methanol, 1:1). Dried samples were dissolved in Mobile Phase B for HPLC.

### Determination of kinetic constants

Kinetic constants were calculated from the Michaelis-Menten representations of at least three ACER3 activity assays, using different concentrations of NBD-C_12_-PHC (from 2.5 to 400 μM), and in the presence of protein amounts from 1 to 5 μg.

### Protein expression analysis

Protein expression was assessed by Western blotting analyses using an anti-FLAG antibody (Sigma-Aldrich, St. Louis, MO) and anti-rabbit IgG (Cell Signaling Technology, Inc., Danvers, MA) secondary antibody.

## RESULTS

To understand the catalytic mechanism of the CREST family of lipid hydrolases, it is important to relate the role of key active site residues with structural information. Alkaline ceramidases, which catalyze the hydrolysis of the amide bond of ceramide to form sphingosine and free fatty acid ^2^, were the first members of CREST superfamily to be identified as lipid hydrolases. However, currently there is no available structural information for the alkaline ceramidases, which has hindered our understanding of their catalytic mechanism. It was previously demonstrated using a bioinformatics approach that ADPRs are related to alkaline ceramidases ^19–20^ and belong to the CREST superfamily of lipid hydrolases ^7^. Recently, the crystal structures of human ADPR1 and ADPR2 were determined. The ADPR structures revealed a 7- transmembrane architecture with a canonical Zn-dependent hydrolase active site located at the bottom of a central cavity ^20^. While the ADPRs have been suggested to be lipid hydrolases, a robust biochemical assay to measure their activity is currently lacking. To determine the catalytic mechanism of the CREST superfamily, we sought to combine the structural information available for the ADPRs, with the biochemical assays available for alkaline ceramidases. We chose to focus our study on ACER3 due to its robust ability to hydrolyze the fluorescent lipid substrate NBD-C_12_-PHC, which can be easily quantitated, and the availability of a genetic mutation that could be used to verify structural models.

### Model of human alkaline ceramidase 3 (ACER3)

To gain insight into the structure and function of alkaline ceramidases, a model of the ACER3 structure was generated using the Phyre2 webserver (http://www.sbg.bio.ic.ac.uk/∼phyre2) based on the crystal structure of ADPR2 ^20^ (Fig. 1). Multiple sequence alignment was performed with the amino acid sequences of ACER1, ACER2, ACER3, ADPR1 and ADPR2 for comparison (Fig. 1D). The majority of the model was predicted with high confidence with only small portions of the protein that corresponded to sites away from the active site predicted with lower confidence.

**Figure 1.**
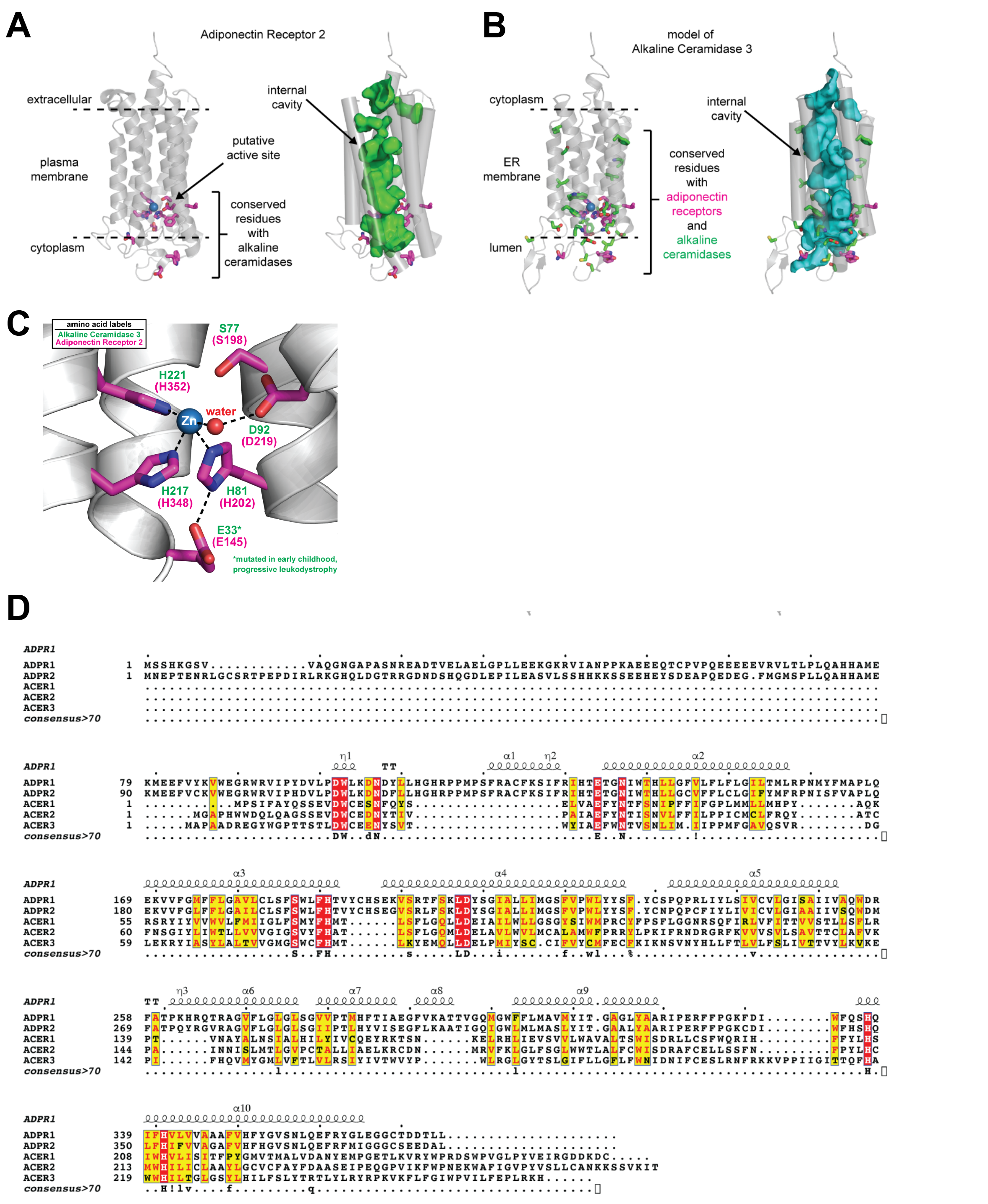
Structure of ADPR2 and modeled structure of ACER3. (A) The structure of ADPR2. Conserved residues with ACER3 in the active site are colored magenta. (B) Modeled structure of ACER3. Conserved residues with ADPRs are colored magenta. Conserved residues with other ACERs (ACER1 and ACER2) are indicated as green. (C) Predicted ACER3 active site highlighting the Zn-coordinating side chains. Corresponding residues in ADPR2 are indicated. (D) Multiple amino acid sequence alignment of ACER1, ACER2, ACER3, ADPR1 and ADPR2. The amino acids conserved in all proteins are boxed by red.

The ACER3 model consisted of a 7-transmembrane architecture with a central hydrophobic cavity similar to ADPRs. The twelve amino acids that are conserved with ADPRs were all clustered on the luminal side of ACER3, either present in the hydrophobic membrane-spanning segment or solvent exposed to the lumen. This included the key catalytic residues consisting of the three His residues involved in Zn-coordination, the Asp residue that hydrogen bonds to the Zn-bound water molecule, and the Ser residue of unknown function that is universally conserved in the CREST superfamily (Fig. 1D). Interestingly, Glu33 was predicted to hydrogen bond to the Zn-coordinating His217, which suggested that this residue is critical for stabilizing the active site (Fig. 1C). This is consistent with the recently described point mutantion E33G, which inactivates ACER3 and causes a rare form of early childhood progressive leukodystrophy ^5^. The remaining conserved residues were either clustered around the core catalytic residues and involved in transmembrane helix-helix stabilizing interactions or were solvent exposed in the lumen.

We also analyzed the position of the residues absolutely conserved among alkaline ceramidases, but not conserved with ADPRs. The majority of these residues were clustered around the key catalytic residues within the active site, which suggests they may play a role in binding and recognizing ceramide. Interestingly, there were only a few residues conserved within the upper portion of the hydrophobic cavity. The general lack of conservation in this region may be due to the specific preference of each ACER for different fatty-acyl chain lengths of ceramide, which is predicted to bind within this hydrophobic cavity. Overall, our model of ACER3 suggests that essential amino acids in the catalytic site are well conserved between ACER3 and ADPRs and suggests that ACER3 is a Zn^2+^-dependent amidase.

### Kinetic Characterization of wild-type and mutated ACER3

To investigate the kinetics of ACER3 ceramidase activity, we used a previously established microsomal assay in conjunction with a newly established HPLC assay with improved sensitivity. First, microsomes were purified from *S. cerevisiae* cells (∆*Ydc1*∆*Ypc1*) overexpressing human ACER3. Since *S. cerevisiae* has only two ceramidases, Ypc1p and Ydc1p, Δ*Ypc1*Δ*Ydc1* cells lack all endogenous ceramidase activity and thus provide the ideal signal to noise ratio to analyze ceramidase activity ^2, 28^. In accordance with previous studies ^2, 18^, the ceramidase activity of ACER3 toward NBD-C_12_-PHC was determined at pH 9.4, unless otherwise mentioned. Prior to determining the enzyme kinetics of ACER3, the linear detection limit of the product NBD-fatty acid was determined (Fig. S1). This newly established method was extremely sensitive and could quantitate NBD-FA levels at far lower levels than our previous TLC based method. In addition, the time and protein concentration for the reaction were optimized and confirmed to be in the linear range (Fig. S2). A plot of the reaction velocity (V) as a function of the substrate concentration (S) obeyed Michaelis-Menten kinetics (R^2^=0.98, Fig. 2A). The *K_M_* and *V_MAX_* values for NBD-C_12_-PHC were calculated to be 15.48 ± 1.248 μM and 46.94 ± 0.8976 pmol/min/mg, respectively (Fig. 2A).

**Figure 2.**
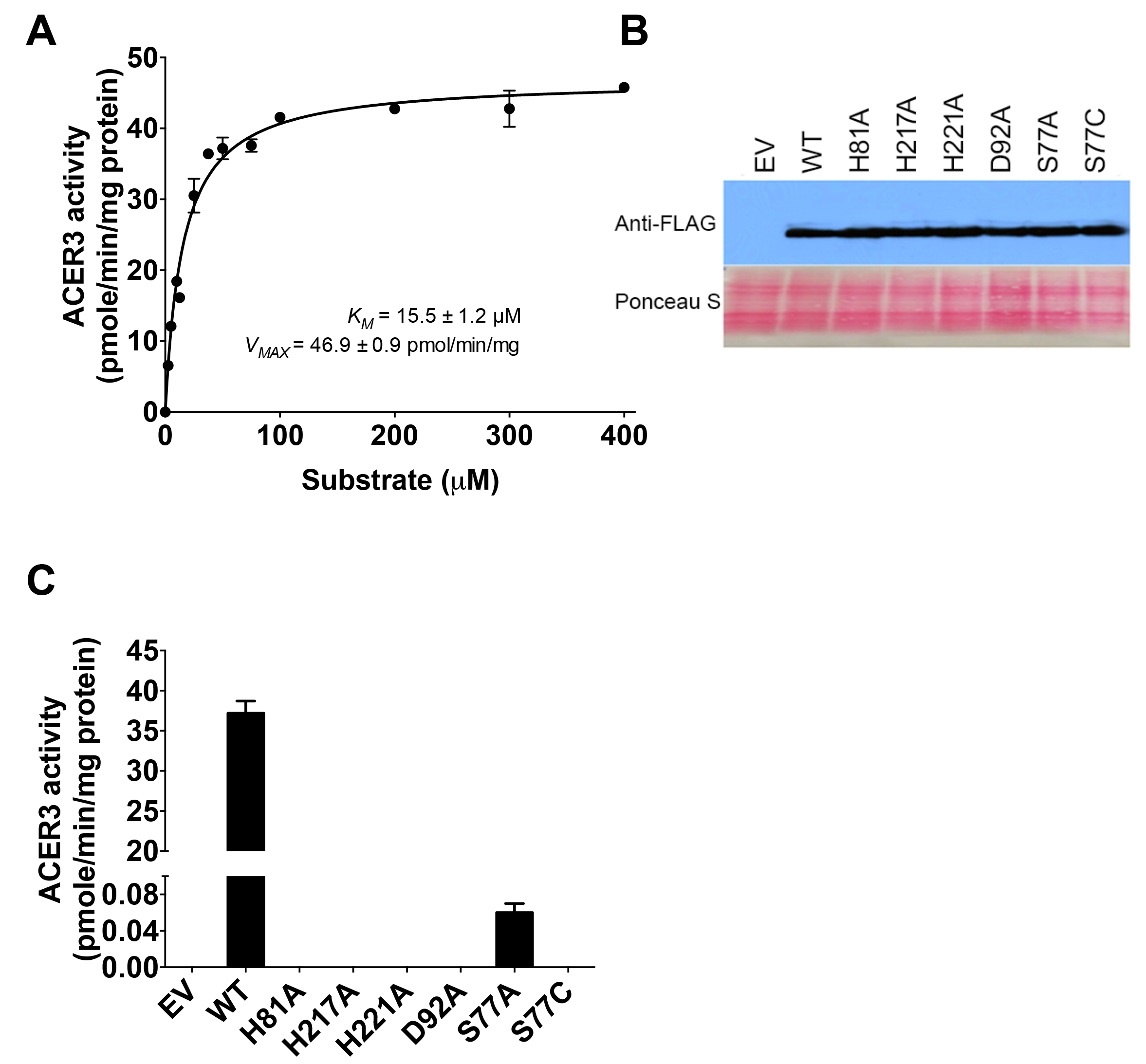
The kinetics of wild-type and mutated ACER3. (A) Microsomes from yeast cells (*ΔYpc1ΔYdc1*) overexpressing wild-type ACER3 were measured for in vitro ceramidase activity on various concentrations of NBD-C12-PHC. *K_M_* and *V_MAX_* values are indicated. Reactions were conducted with 1 μg of microsomal proteins at 37 °C for 30 min. Data represent the mean ± S.D. of three independent experiments performed in duplicate. (B) Western blot analysis was performed with the anti-FLAG antibody to verify the expression levels of each mutated and wild-type ACER3. For normalization of equal loading, the membrane blot was stained by ponceau S. (C) Microsomes from wild-type and each mutated ACER3 were subjected to alkaline ceramidase activity using NBD-C_12_-PHC. The release of the fluorescent product NBD-C_12_-FA from the substrate NBD-C_12_-PHC was detected by HPLC. EV, empty vector.

Our model of ACER3 identified a canonical Zn-amidase active site that included three Zn-coordinating His residues and an Asp residue that was hydrogen-bonding with a Zn-bound water. To test the hypothesis that ACER3 is a Zn-dependent amidase, we investigated the importance of each active residue of ACER3 by site-directed mutagenesis. Each residue was replaced with Ala to make the point mutations of ACER3 including H81A, H217A, H221A, and D92A. The expression levels of wild-type and all mutated ACER3 were verified to be similar by western blot (Fig. 2B). Ceramidase activities of wild-type and the four point mutations toward NBD-C_12_-PHC were measured by HPLC (Fig. 2C). Ceramidase activity was not detected in all of the mutations. This suggested these residues were all essential for ACER3 to catalyze the hydrolysis of NBD-C12-PHC.

### Mutational analysis of Ser77

Unlike other Zn-dependent amidases, ACER3 contains a unique serine residue located near the Zn^2+^ ion in the active site. This Ser residue is universally conserved in ADPRs and the CREST superfamily and is part of the motif that defines this protein superfamily. However, the role of S77 is unclear as other Zn-dependent amidases do not contain a Ser residue in their active site. To ascertain the role of Ser77, we used site-direct mutagenesis to replace Ser77 with Ala (S77A) and determined the effect on ACER3 ceramidase activity. Interestingly, S77A had a low but measurable ceramidase activity that was approximately 600-fold lower than WT ACER3 (Fig. 2C). The reaction of S77A followed Michaelis– Menten kinetics with a dramatically reduced *V_max_* (0.061 vs. 47 pmol/min/mg) that was accompanied by a slight increase in *K_M_* from 15.5 μM to 38 μM (Fig. 3A).

**Figure 3.**
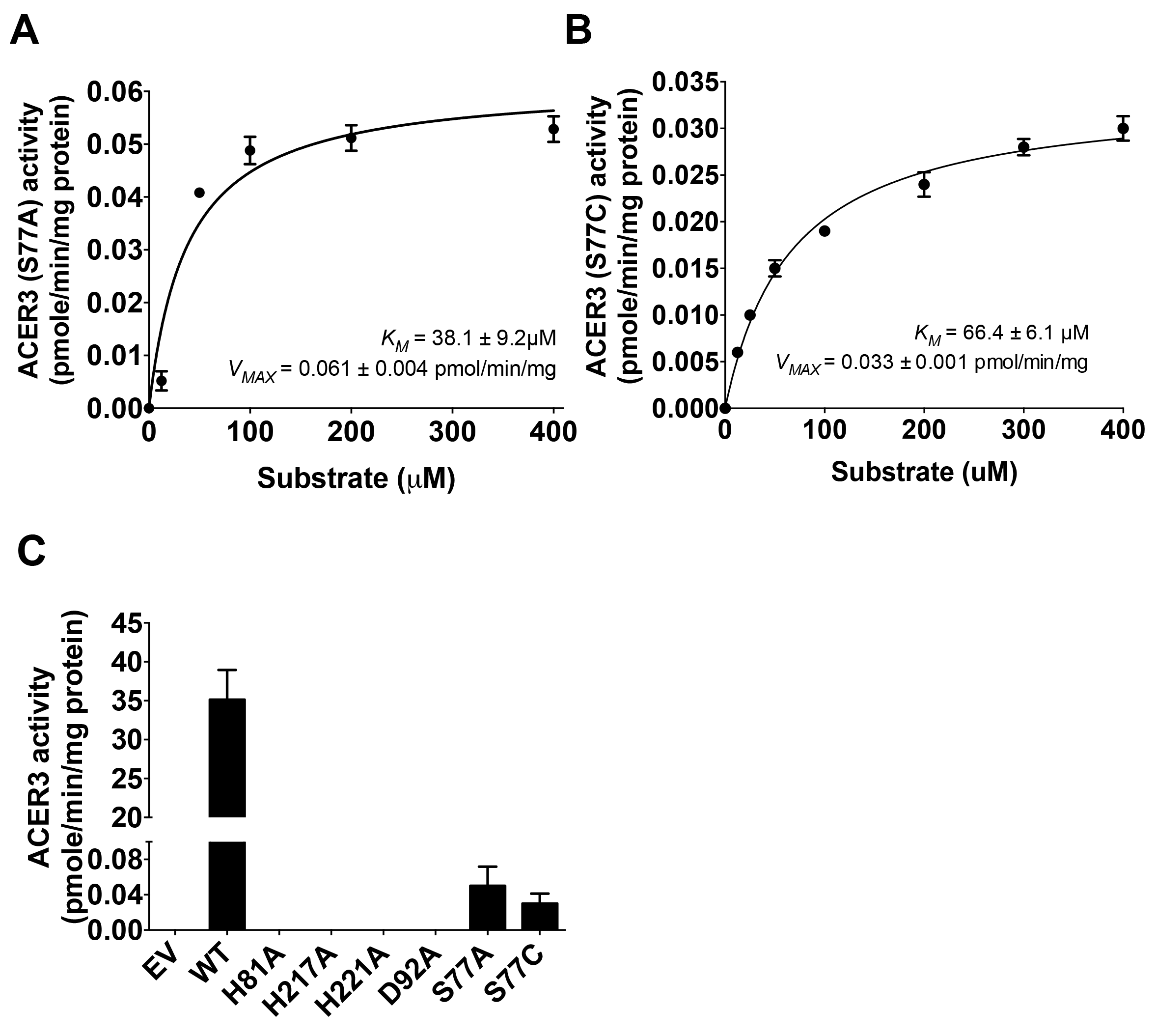
Role of Ser77 in ACER3 catalytic site. (A) Microsomes from yeast cells (*ΔYpc1ΔYdc1*) overexpressing mutated ACER3 (S77A) were measured for in vitro ceramidase activity on various concentrations of NBD-C12-PHC. *K_M_* and *V_MAX_* values are indicated. Reactions were conducted with 1 μg of microsomal proteins at 37 °C for 30 min. Data represent the mean ± S.D. of three independent experiments performed in duplicate. (A) Microsomes from WT and each mutated ACER3 were subjected to alkaline ceramidase activity using NBD-C_12_-PHC at pH 7.5. The release of the fluorescent product NBD-C_12_-FA from the substrate NBD-C_12_-PHC was detected by HPLC. (C) Microsomes from S77C were measured for in vitro ceramidase activity on various concentrations of NBD-C_12_-PHC at pH 7.5. *K_M_* and *V_MAX_* values are indicated. Reactions were conducted with 1 μg of microsomal protein at 37 °C for 30 min. Data represent the mean ± S.D. of three independent experiments performed in duplicate. EV, empty vector.

To further investigate the role of Ser77, we generated the point mutation S77C using site-directed mutagenesis. This point mutation was chosen because we reasoned that the thiol group (- SH) of cysteine might be able to complement the hydroxyl group (-OH) of serine. However, the S77C ACER3 point mutation had no measurable alkaline ceramidase activity. Since the pKa of the cysteine thiol group is 8.14 and we conducted our analysis at the ACER3 pH optima of 9.4. we evaluated if lowering the pH to 7.5, which would protonate the thiol group and eliminate the negative charge, would be sufficient to restore some ACER3 activity (Fig. 3B and 3C). Interestingly, decreasing the pH to 7.5 showed a notable increase of S77C activity, equivalent to the S77A mutation. At pH 7.5, wild-type ACER3 had a slight 7% reduction of activity and there were no measurable effects on all other mutations (Fig. 3C). Since the recovered activity of S77C was very low, we measured the enzyme kinetics of S77C to validate if the recovered activity of S77C comes from ACER3 enzyme reaction (Fig. 3B). The *K_M_* and *V_MAX_* values were computed to be 66.43 ± 6.143 μM and 0.0337 ± 0.01 pmol/min/mg, respectively, showing that the plot obeys Michaelis-Menten kinetics. These results suggest that the thiol group of cysteines partially complement the hydroxyl group (-OH) of serine.

### Pharmacological inhibition of ACER3

There has been an effort to seek specific inhibitors of alkaline ceramidases. Additionally, HDAC (class I/II) inhibitors are well known to interrupt Zn^2+^- dependent amidases ^29^. We decided to test if these inhibitors have an effect on ACER3 ceramidase activity. Microsomes from *S. cerevisiae* cells overexpressing wild-type ACER3 were treated with inhibitors including trichostatin A (TSA) or C6-Urea-Ceramide, which is a neutral ceramidase inhibitor ^30^. C6-Urea-Ceramide did not have an effect on ACER3 activity at concentrations up to 30 μM (Fig. 4A). Only TSA treatment inhibited the activity in a concentration dependent manner. To determine the mechanism of inhibition of TSA, we performed kinetic assays. As seen in Figure 4B, TSA treatment had no effect on the *K_M_* (*K_M_* = 15.89 ± 1.077 μM) but reduced the *V_MAX_* (*V_MAX_* = 31.53 ± 0.493 pmol/min/mg) of ACER3. This is indicative of an non-competitive mechanism of inhibition (Fig. 4B).

**Figure 4.**
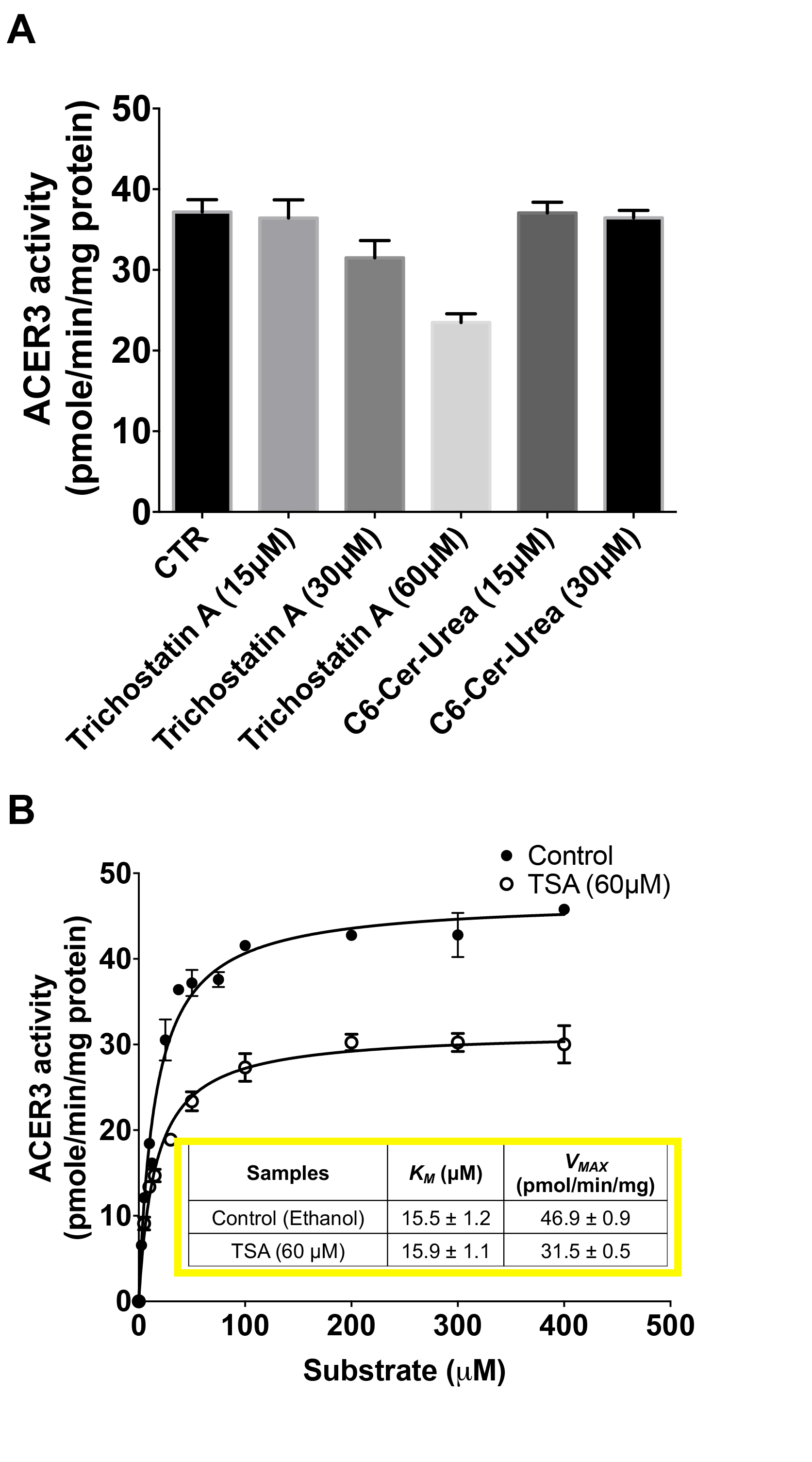
Inhibitor assay of ACER3. (A) Microsomes from yeast cells (*ΔYpc1ΔYdc1*) overexpressing wild-type ACER3 were collected and treated with different inhibitors (TSA and C6-Cer-Urea). In turn, they were subjected to alkaline ceramidase activity using NBD-C_12_-PHC. The release of the fluorescent product NBD-C_12_-FA from the substrate NBD-C_12_-PHC was detected by HPLC. CTR, non-treated. (B) Microsomes treated with TSA (60μM) were measured for in vitro ceramidase activity on various concentrations of NBD-C_12_-PHC. The reaction was plotted against concentration and merged with control for the quick comparison. The *K_M_* and *V_MAX_* values are indicated. Data represent the mean ± S.D. of three independent experiments performed in duplicate.

## DISCUSSION

Based on the predicted model of ACER3 and our mutational analysis, we propose a general acid-base catalysis mechanism for ceramide hydrolysis by ACER3. By our suggested mechanism, Asp92 is predicted to activate the water molecule by deprotonation, serving as a general base (Fig. 5). The activated water molecule undergoes a nucleophilic attack on the amide bond resulting in the formation of an oxyanion bound to a tetrahedral carbon. The terminal amine of the sphingosine moiety is predicted to abstract the proton from Asp92, resulting in the collapse of the tetrahedral carbon intermediate, releasing sphingosine and a fatty acid. Finally, the protein catalyst is regenerated.

**Figure 5.**
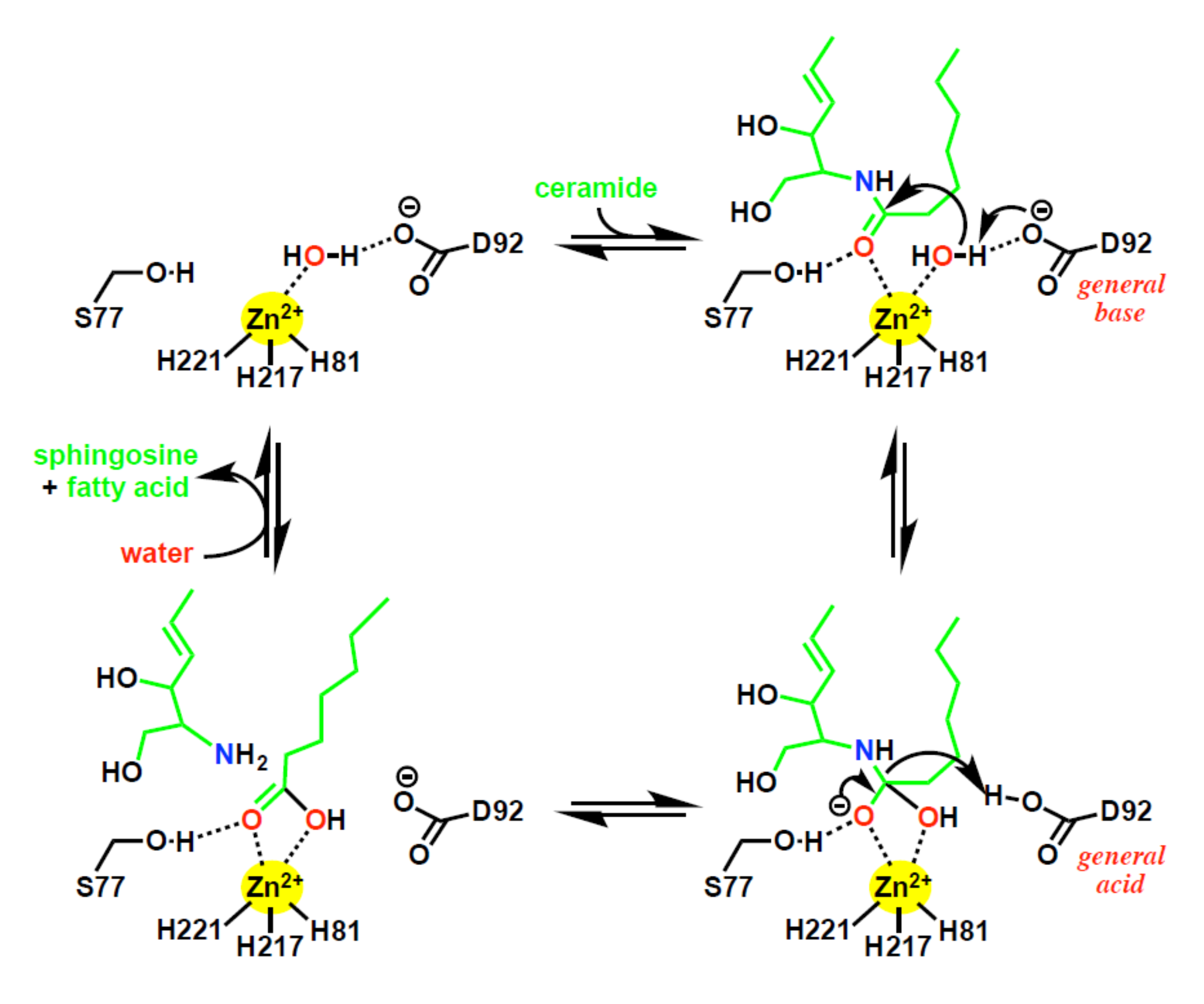
Suggested ACER3 catalytic mechanism. The bottom right panel, tetrahedral transition state.

Consistent with this mechanism, the mutations of the three proximal Zn-coordinating residues (His81, His217, and His221) and Asp92 abolished ACER3 activity (Fig. 2C). Additionally, substitutions of Ser77 dramatically decreased ACER3 ceramidase activity suggesting the important role of Ser77. Ser77 is predicted to form a hydrogen bond with the oxygen of carbonyl from the amide bond of ceramide, to help position the amide bond of ceramide for nucleophilic attack by the water molecule. Ser77 is also predicted to stabilize the negatively charged oxyanion of the transition state. The proposed role of Ser77 is supported by the pH dependence of the S77C mutation (Fig. 3B and 3C). At alkaline pH the cysteine residue is unprotonated and negatively charged, thus it would be unable to hydrogen bond with the oxygen carbonyl and would interact unfavorably with the negatively charged oxyanion. Overall, our proposed mechanism identifies key roles for all the universally conserved residues of the CREST superfamily in the active site of ACER3.

Importantly, our proposed mechanism does not involve rearrangement of the Zinc-coordinating residues as proposed for ADPRs ^19^. In the proposed ADPR mechanism, the conserved Ser residue is proposed to be involved in ceramide hydrolysis. To be specific, they suggest that the rearrangement of the zinc binding site upon ceramide binding can lead to the direct coordination of Ser77 hydroxyl to the zinc. However, the low, but the measurable activity of the S77A and S77C mutations is not consistent with a role for the conserved Ser in metal coordination. Based on the pH sensitivity, it appears it is most likely involved in aiding stabilization of the transition state oxyanion. In support of this, a similarly positioned Threonine residue in LpxC has been proposed to play a similar role in stabilizing the oxyanion of the transition state ^31^.

Lastly, as ACERs have emerged as therapeutic targets for cancer, this study provides important insight into the type of inhibitors that may be suitable for targeting ACERs. There are several therapeutic targets, including HDACs, LpxC, peptide deformylase, MMPs, and neutral ceramidase that have similar active sites. Hydroxamate compounds, such as TSA, have proven effective inhibitors for this class of enzymes. Especially, it has been reported that TSA treatment leads to increase in endogenous ceramides in human cancer cells ^32^. In turn, increased ceramides induced cell death, suggesting a potential inhibitory effect on ACERs by TSA in human cells as well as in test tubes. Given the similar Zn^2+^-based active site of ACER3 and the canonical catalytic mechanism we propose, it is likely that hydroxamates, which display strong binding to Zn centers, may be developed as inhibitors of the ACERs for cancer therapy.

## ACKNOWLEDGEMENTS

This work was supported, in whole or in part, by National Institutes of Health Grants R01CA163825 (to C.M), P01CA097132 (to Y.A.H and C.M), GM062887 (to L.M.O) and a grant from the American Heart Association 17SDG33410860 (to M.V.A.)

## CONFLICT OF INTEREST STATEMENT

The authors declare that they have no conflicts of interest with the contents of this article.

---

**Figure S1. HPLC chromatogram of standard NBD-C_12_-FA** (A) HPLC chromatogram of different concentrations of NBD-C_12_-FA (100 fM, 1 pM, 10 pM, and 100 pM) obtained from HPLC-FLD Fluorescent Detector (Agilent, Santa Clara, CA) set to excitation and emission wavelengths of 467 and 540 nm, respectively. (B) Merged HPLC chromatogram of standard NBD-C_12_-FA (C) Calibration curve for NBD-C_12_-FA. Equation: Y = 1.346*X + 2.194, R^2^=0.9999

**Figure S2. Linearity of ACER3 activity assay** (A) Assay linearity with reaction time and three different protein amounts (0.5 μg, 1 μg, and 2 μg). The graph shows NBD-C_12_-FA (pmole) versus the reaction time for each different amount of microsome preparation. (B) Assay linearity with protein amount. The graph shows NBD-C_12_-FA (pmole) versus the amount of microsome for 30 min. Data represent the mean ± S.D. of three independent experiments performed in duplicate.

**Figure S2. HPLC chromatogram of ACER3 reaction** Upper panel shows HPLC chromatogram of wild-type ACER3 reaction. The lower panel shows HPLC chromatogram of empty vector reaction. The substrate (NBD-C_12_-PHC) and product (NBD-C_12_-FA) are indicated. Reactions were conducted with 1 μg of microsomal proteins at 37 °C for 30 min.

